# Sea of Electrodes Array (SEA): Extremely Dense and High-Count Silicon-Based Electrode Array Technology for High-Resolution High-Bandwidth Interfacing with 3D Neural Structures

**DOI:** 10.1101/2021.01.24.427975

**Authors:** Amin Sandoughsaz Zardini, Behnoush Rostami, Khalil Najafi, Vaughn L. Hetrick, Omar J. Ahmed

## Abstract

In this work, we propose a new silicon-based micro-fabrication technology to fabricate 3D high-density high-electrode-count neural micro-probe arrays scalable to thousands and even millions of individual electrodes with user-defined length, width, shape, and tip profile. This unique technology utilizes DRIE of ultra-high aspect-ratio holes in silicon and refilling them with multiple films to form thousands of individual needles with metal tips making up the “sea-of-electrodes” array (SEA). World-record density of 400 electrodes/mm^2^ in a 5184-needle array is achieved. The needles are ~0.5-1.2mm long, <20μm wide at the base, and <1μm at the tip. The silicon-based structure of these 3D array probes with sharp tips, makes them stiff enough and easily implantable in the brain to reach a targeted region without failing. Moreover, the high aspect ratio of these extremely fine needles reduces the tissue damage and improves the chronic stability. Functionality of the electrodes is investigated using acute *in vivo* recording in a rat barrel field cortex under isoflurane anesthesia.

## Introduction

The vast endeavors of neuroscientists for more than two centuries have led to a partial understanding of the mechanism of the brain activities through the advances in neuronal technologies. To date, different technologies have been developed for brain monitoring such as EEG, fMRI, ECoG [1], and implantable neural interfaces through modulating and detecting the local field potentials (LFP) [2] and single unit spikes [3]. However, decoding the neural circuits down to a single neuron level is solely enabled through implantable neural probes [4].

There is a long history of research for developing intracortical neural probes comprising individual microelectrodes for large-scale recording of neuronal activities [5–7]. The continuous tendency toward studying larger number of neurons has led to doubling of simultaneously recorded neurons every 7 years since the 1950’s [8]. Conventionally, neural probes are classified based on their geometry into two types of 2D planar probes and 3D out-of-plane arrays. In planar probes, recording sites are laid out along a single shank and can cover neighboring neurons in 1D to 1.5D fashions as shown in Fig.1 (A) and (B). The recording plane in planar probes is primarily perpendicular to the surface of the cortex, i.e., the recording plane is the same as the plane formed by the probe shanks. Michigan probe is one of the most popular examples of planar probes with 1D recording sites coverage which has been extremely useful for many years. Obtaining large number of recording sites in planar probes while preserving minimally invasiveness, is mostly limited by the shank size. In other words, large number of electrodes along a single shank is obtained at the expense of larger shank size which will cause tissue damage and reduces chronic stability. Neuralink probes with 32 sites per thread is an example of the latest polymer-based planar probes with 1D-1.5D spatial coverage [9]. Another example is the IMEC silicon-based probe with 1600 sites on a single shank with 1.5D sites spatial coverage as one of the probes with highest site count per shank [10]. Multiple-shank architecture has been used to scale up the site counts without increasing the probe shank cross-sectional footprint. Neuralink 96-thread array with 3072 [9] sites and IMEC 4-shank with 5120 sites [11] are the latest probes of this type.

**Figure1.**
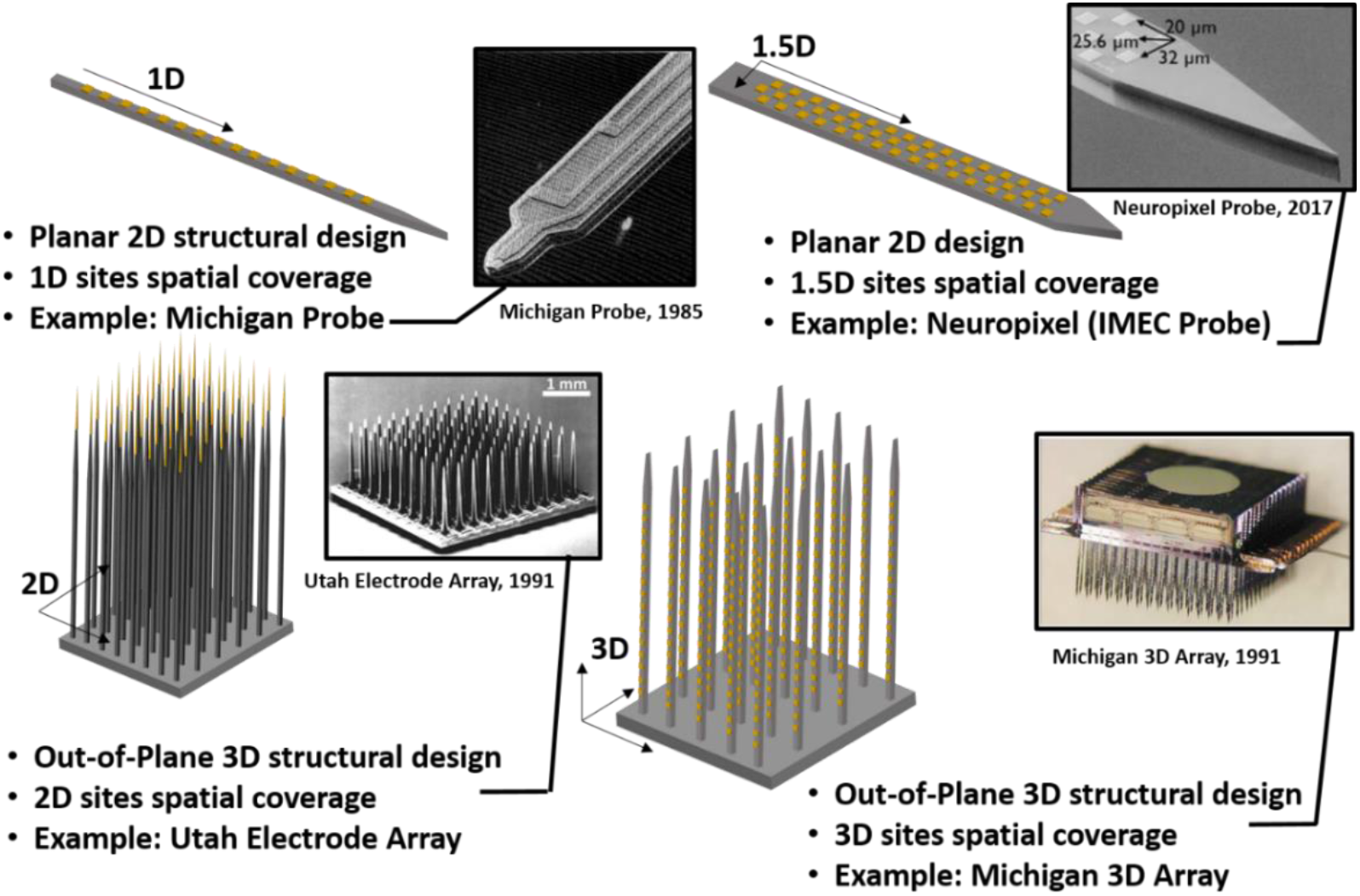
Electrode arrays structural design and spatial coverage: (A) planar probes with linear 1D sites spatial coverage, (B) planar probes with increased spatial coverage along the width of the shank classified as 1.5D spatial coverage, (C) out-of-plane architecture with 2D planar spatial coverage, (D) 3D out-of-plane architecture with true 3D volume spatial coverage.

Due to the invasiveness concerns, increasing the probe dimensions is limited. Therefore, there is not much space for increasing the number of sites on a planar probe. 2D probes can alternatively be formed by an array of electrodes whose tips form the recording sites, which form a recording plane as shown in Fig.1 (C). In this case, the recording plane is primarily parallel to the cortical surface, allowing recording from a single neural layer. Utah Electrode Array (UEA) with 96 sites is the most well-known probe of this kind [12]. 2D arrays have also been demonstrated with significant improved capabilities and with many more sites by bundling insulated microwires or fibers [13–15], or by assembling carbon fibers [16–19], each of which forms a recording site at the tip. The Utah probe, like the microwire or carbon bundles, is 3D in structure but primarily 2D in its recording ability. The slanted Utah array [20] allows recording from multiple neural layers with its recording plane angled to the cortical surface and is referred to as a 2.5D probe. In case of bundled wires/fibers probes, the location, exact distribution, and footprints of the sites are obviously not controllable after the tissue insertion.

Most of the existing 1D and 2D probes exhibit excellent in-depth spatial resolution; however, their poor areal coverage due to their large shank size or large gap between the shanks, limits their use in applications which require large-scale recording within a specific depth of brain. In this case, probes with wider shanks or multiple parallel probes are needed to cover a larger area which both will result in more tissue damage and less chronic stability. On the other hand, out-of-plane electrode arrays can potentially provide sufficient planar coverage; however, their depth coverage is still poor.

These shortcomings, in terms of latitudinal and longitudinal spatial resolution, have hindered true three-dimensional neural studies with high spatial resolution. Moreover, complex geometry of the brain includes some non-planar regions that are hard to reach by planar probes. To interface three dimensionally with the neuronal populations, different types of 3D out-of-plane arrays of microelectrodes have been developed. An example of 3D arrays is shown in Fig.1 (D) in which multiple sites are distributed along the length of several shanks, which are themselves distributed along an area of tissue. The Michigan 3D probe reported is the earliest 3D probe array of this kind for both recording and stimulation [21–23]. Rios at Caltech [24] has also demonstrated a large-count 3D probe by stacking several individual planar multi-shank and multi-site silicon probes. However, Michigan and Caltech 3D arrays are still limited in channel count due to their large footprints that causes high volumetric displacement and tissue damage [25].

Through all these years of studies, researchers have been attempting to develop an ideal neural technology to modulate and detect neuronal activities simultaneously from large-scale arbitrary distributions within the 3D neuronal populations while hindering brain tissue damages. Despite many research activities, we are still some way from building an “ideal” probe. In fact, the “ideal” probe for mapping the activities of individual neurons and neural circuits might not be in reach for many more decades because the ideal probe needs to record from many neurons through a 3D volume of tissue and be able to achieve this with no or minimal damage over a long period of time. However, we can come close to this ideal probe, and possibly produce a “near-ideal” probe with features that help overcome the shortcomings of existing probes. Fig. 2 is an attempt to conceptually illustrate these near-ideal probe features and provide a roadmap for the future transformative neural technologies. These features are briefly discussed through an example of recording from a rat brain’s hippocampus. Hippocampus has one of the highest density of neurons which is extended through various layers of brain, therefore a near-ideal probe requires to provide these features:

1. **Large-scale high-resolution recording:** high-count dense electrode arrays to access a large population of neurons with high dexterity are essential for accurate mapping of the brain, Fig. 2(A).
2. **3D spatial coverage:** to access neurons in various depth and span of brain structures with 3D irregular shapes such as hippocampus ram’s horn shape structure. Ideally with precise control over the position and placement of recording/stimulation sites following implantation, Fig. 2(A).
3. **Material and structural robustness:** sufficient mechanical strength for successful insertion and durability post-implantation, material biocompatibility to prevent cell toxicity or triggering the cellular immune reactions and chronic *in vivo* stability of the electrodes in tissue harsh environment to realize long-term stable access to neurons in particular for clinical applications. These requirements also apply to the probe encapsulation, interconnections and packaging, Fig. 2(B).
4. **Minimal invasiveness:** minimizing the tissue volume displacement and tissue damage by reducing the electrodes dimensions, increasing the tip sharpness, and reducing the mismatch between brain and probe’s Young’s modulus, Fig. 2(B).
5. **Multi-modality:** ability to record/stimulate electrically, and interface optically and chemically through a single shank, Fig 2(C-E).

**Figure 2.**
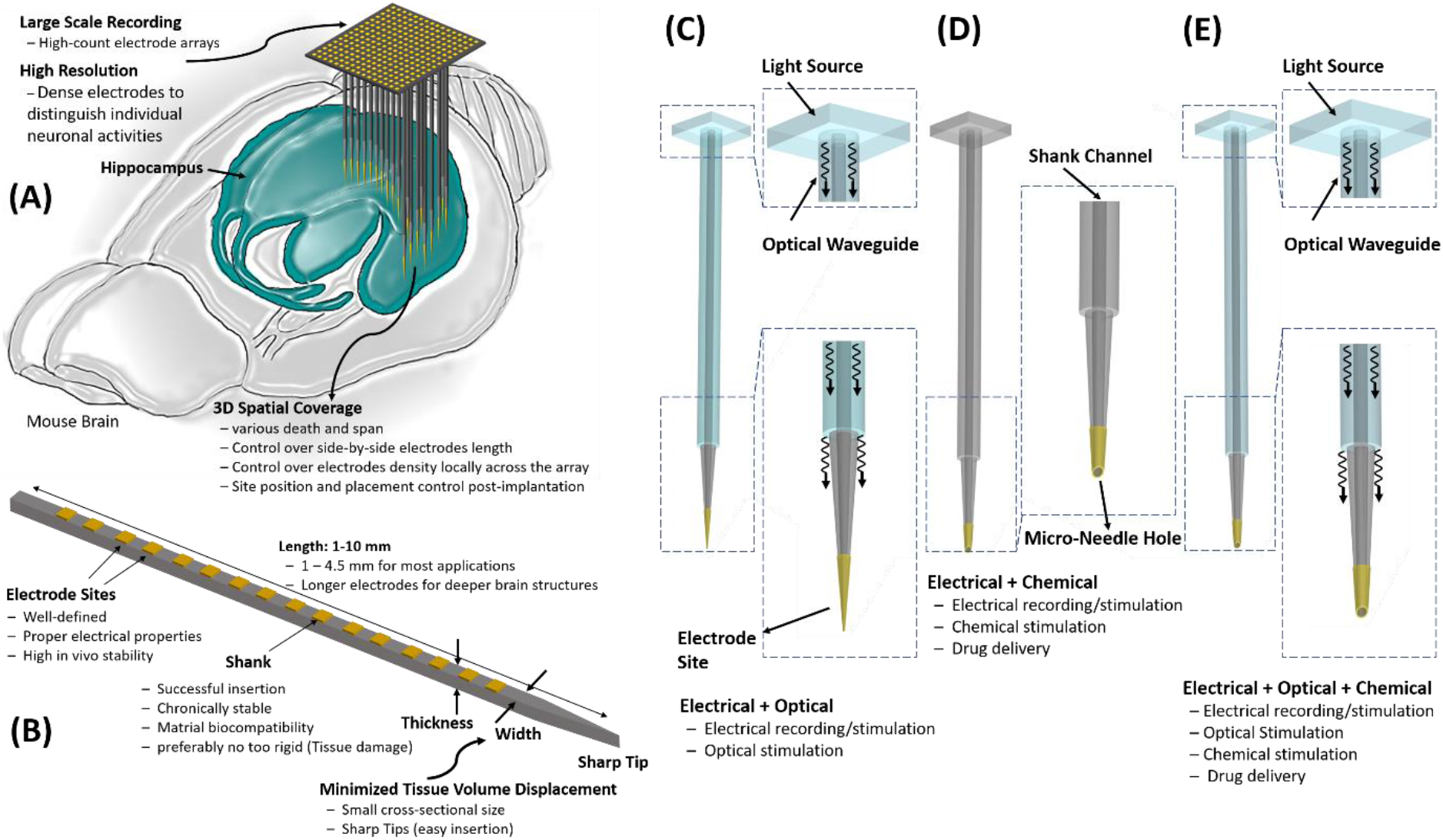
Conceptual illustration of a near-ideal probe features: (A) large scale high resolution electrode array with true 3D spatial coverage to interface with neurons in various depth/span with control over site position and placement post-implantation, (B) material and structural requirements: sufficient robustness for successful insertion and survival post-implantation, material biocompatibility and chronic stability, electrode sites with proper electrical properties and *in vivo* stability, Minimized invasiveness: sharp tips and small cross-sectional footprint to reduce tissue damage, tissue volume displacement and minimizing the rigidity mismatch between tissue and probe shanks, (C) electrophysiological and optical modalities, (D) electrophysiological and chemical modalities, (E) electrophysiological, optical and chemical modalities provided in a single probe shank for electrical, optical and chemical interfacing with neurons.

Here, we present a novel technology capable of manufacturing electrode arrays with site counts well beyond the state-of-the-art projection, providing neuroscientist the near ideal probes they have always been deprived due to technological barrier.

## Results and Discussion

The SEA manufacturing process employs a scalable technology to fabricate a new generation of 3D arrays with high-density high-electrode-count neural recording electrodes made from silicon. To overcome the shortcomings and issues of previously reported arrays, a fabrication process based on refilling deep ultra-high aspect-ratio holes in a silicon substrate with deposited layers followed by etching away the support substrate to leave thousands and eventually millions of electrodes, is developed.

Fig. 3 shows an overview of the refilling-based fabrication technology. Details of each step are discussed in the following sub-sections. The fabrication process begins with a custom high aspect-ratio deep reactive ion etching (DRIE) of holes in a p-type *(100)* silicon wafer, Fig. 3(A). DRIE Aspect Ratio Dependent Etching (ARDE) effect and DRIE lag effect are utilized to obtain a tapered profile at the bottom of the holes, which is required to obtain electrodes with conical shape and sharp tips. These holes are then refilled with multiple layers of low-pressure chemical vapor deposited (LPCVD) films to form the electrode conductor and insulator layers. In the current process, we deposit the following layers in order: 1) a sacrificial polysilicon layer (which is etched away in the final structure) of a few microns in thickness to reduce the hole diameter and thus the final needle shank diameter, Fig. 3(B); 2) a composite layer consisting of LPCVD Oxide-Nitride-Oxide (ONO) to form the outer insulator of each individual electrode, Fig. 3(B); 3) a second LPCVD polysilicon layer that is highly doped to form the conductive core of the electrode,

**Figure 3.**
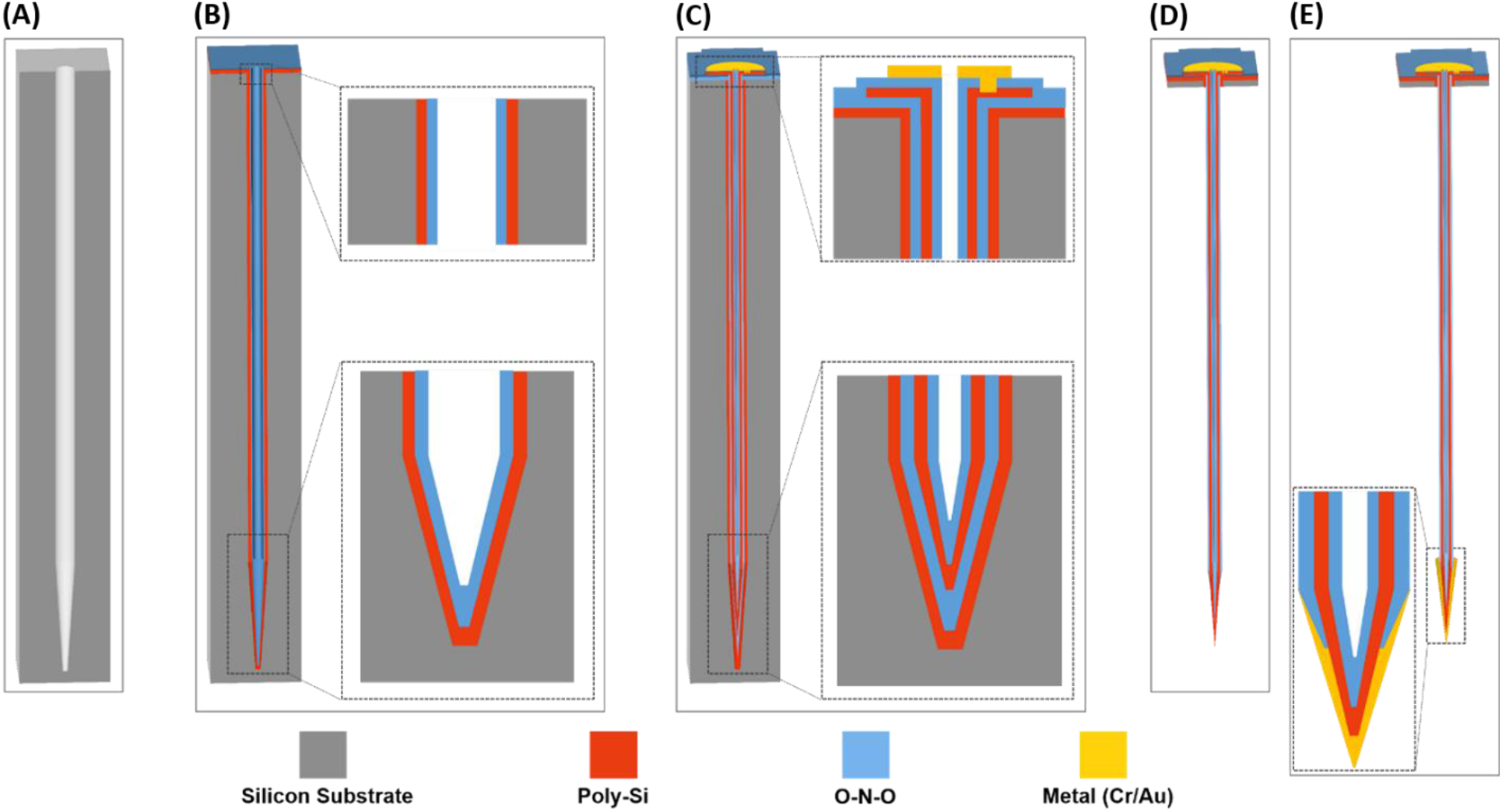
High-density high-electrode-count probe array fabrication technology: (A) Deep ultra-high aspect-ratio DRIE of <25μm diameter holes, DRIE lag effect is utilized to get tapered profile at the bottom of holes which is required to obtain sharp electrode tips. (B) Refilling of holes with multiple low-pressure chemical vapor deposited (LPCVD) films: 1-sacrificial polysilicon to reduce shank thickness, 2-outer oxide-nitride-oxide (ONO) insulator. (C) n-type polysilicon and ONO LPCVD as the electrode conductive core and the inner ONO insulator layer respectively followed by Contact pads patterning on wafer top to electrically access the electrode conductive core. (D) Dissolving the substrate and sacrificial silicon by EDP etching. (E) Maskless RIE etching of the ONO layer at the tip to expose the n-type polysilicon and electrodes tip metallization using gold electroplating process to form ohmic contact to obtain proper impedance required for electrophysiological recording.

Fig. 3(C); and 4) a second LPCVD ONO inner insulator layer, Fig. 3(C). The ONO insulator layers are used for electrical insulation and protection of the n-type polysilicon layer in the following wet etching of the silicon substrate needed to reveal the electrodes. At this point the holes refilling step by LPCVD layers is completed and the top surface of the wafer is flat. The top dielectric ONO layers are removed selectively to provide electrical access to the core polysilicon conductor and a metal layer is deposited in these contact regions to create pads for electrical connection to each of the sites, Fig. 3(C). In Fig. 3 we only show one individual electrode. Needless to say, the process can form hundreds of thousands of these needles side by side. Next, a wet silicon etchant such as EDP is used to dissolve the substrate and sacrificial polysilicon, leaving behind insulated and instrumented electrodes, Fig. 3(D). The next step is to expose the tip and coat it with a conductor to form the recording/stimulation site. This is achieved by doing a mask-less blanket silicon plasma etch to expose the electrodes and then using an oxynitride RIE etch to expose only the tip by etching the outer ONO layer and metallizing the tip, typically with gold, using gold electroplating deposition technique, Fig. 3(E).

In many applications such as brain-computer interfaces (BCI) for restoring limb, full-body movements, restoring vision, hearing and memory, recording/stimulation of higher number of neurons is required, therefore, thousands or even millions of channel counts with high density may be required. The electrode arrays reported to date have a limited channel count (up to 1000), therefore, are not applicable in cases when a dense array with high electrode count is needed.

Using the fabrication process described above, we have fabricated an array with 72×72 electrodes to demonstrate the scalability of our approach in probe site numbers. This array consist of 5184 electrodes with 50μm pitch, each electrode being ~500μm long, 20μm thick at the base and <2μm at the tip as shown in Fig. 4(A-B). Our technology can be readily employed to realize arrays with thousands and even millions of electrodes. Fig. 4(C) shows a single electrode after etching of ONO layers at the tip and exposing the tip n-type polysilicon. Next, gold electroplating process will be used to metallize the electrode tips and form the ohmic contacts, i.e., the recording sites as shown in Fig. 4(D).

**Figure 4.**
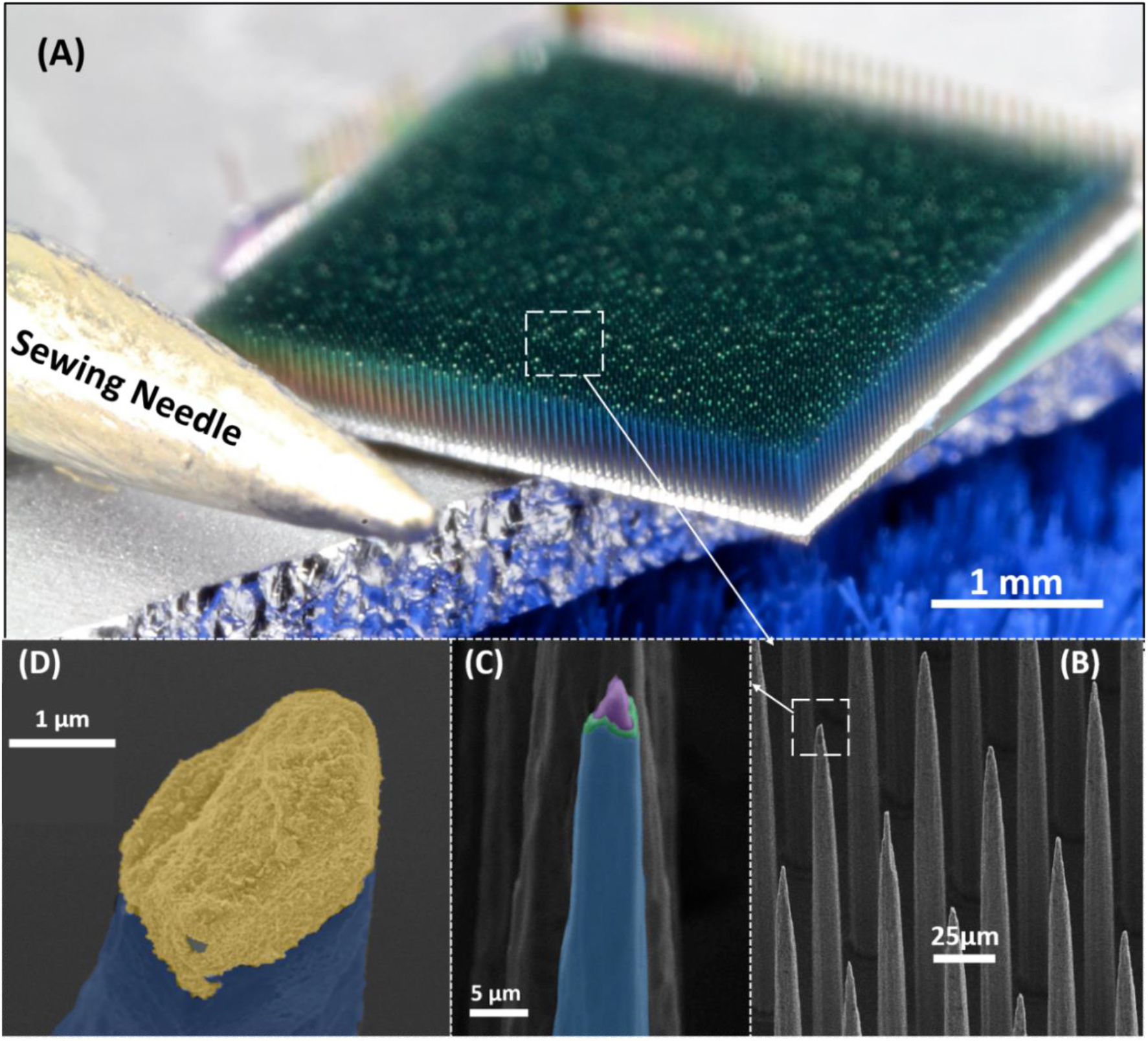
Demonstration of scalability of the SEA fabrication process: (A) 72×72 electrode array with 5184 electrodes on a 3.6×3.6 mm^2^ footprint next to a sewing needle to show the fine dexterity of the array. Each electrode is ~500μm long, 20μm thick at the thickest base and <2μm at the tip with a tapered profile towards the tip which facilitates the tissue insertion. (B) Close-up view of electrodes in a SEM image, (C) False-colored SEM image of a single electrode with exposed conductor layer at the tip. Blue, green and purple represent the electrical insulator SiO2-Si3N4 layers and conductive n-type polysilicon layer, respectively. (D) False-colored SEM image of the recording site after gold-platin the tip. Blue and gold colors represent the insulating SiO2 and gold-plated tip, respectively.

### Modified Process to Obtain Millimeter-long Electrodes

Human Brain neural recording/stimulation applications require electrodes with lengths of 1 to 100 mm [26], while centimeter-long probes are needed mainly for reaching deep structures in human subjects, millimeter-long probes are needed even for non-human subject. In the fabrication process presented in Fig. 3, the electrode length is defined by the DRIE Aspect-Ratio Dependent Etch (ARDE) and DRIE lag effects [27], therefore, for opening holes with a diameter of <30 μm, the electrode length is limited to ~500 μm. In order to obtain neural arrays with millimeter-long electrodes, we have developed another novel process which is based on bonding of multiple silicon wafers. This approach starts with preparing multiple separate wafers containing through-wafer etched holes using a special ramped-parameter ultra-deep high-aspect-ratio DRIE process (UDRIE) as described in [28]. On the other hand, one wafer is etched using the custom DRIE fixed-parameter recipe also described in [28] to utilize the DRIE lag effect to form the tip shape and sharpness, Fig. 5(A). This and all other wafers are bonded to form mm-deep holes as shown in Fig. 5(B). These long holes are again refilled with the same fabrication process described in Fig. 3 to obtain mm-long electrode arrays.

**Figure 5.**
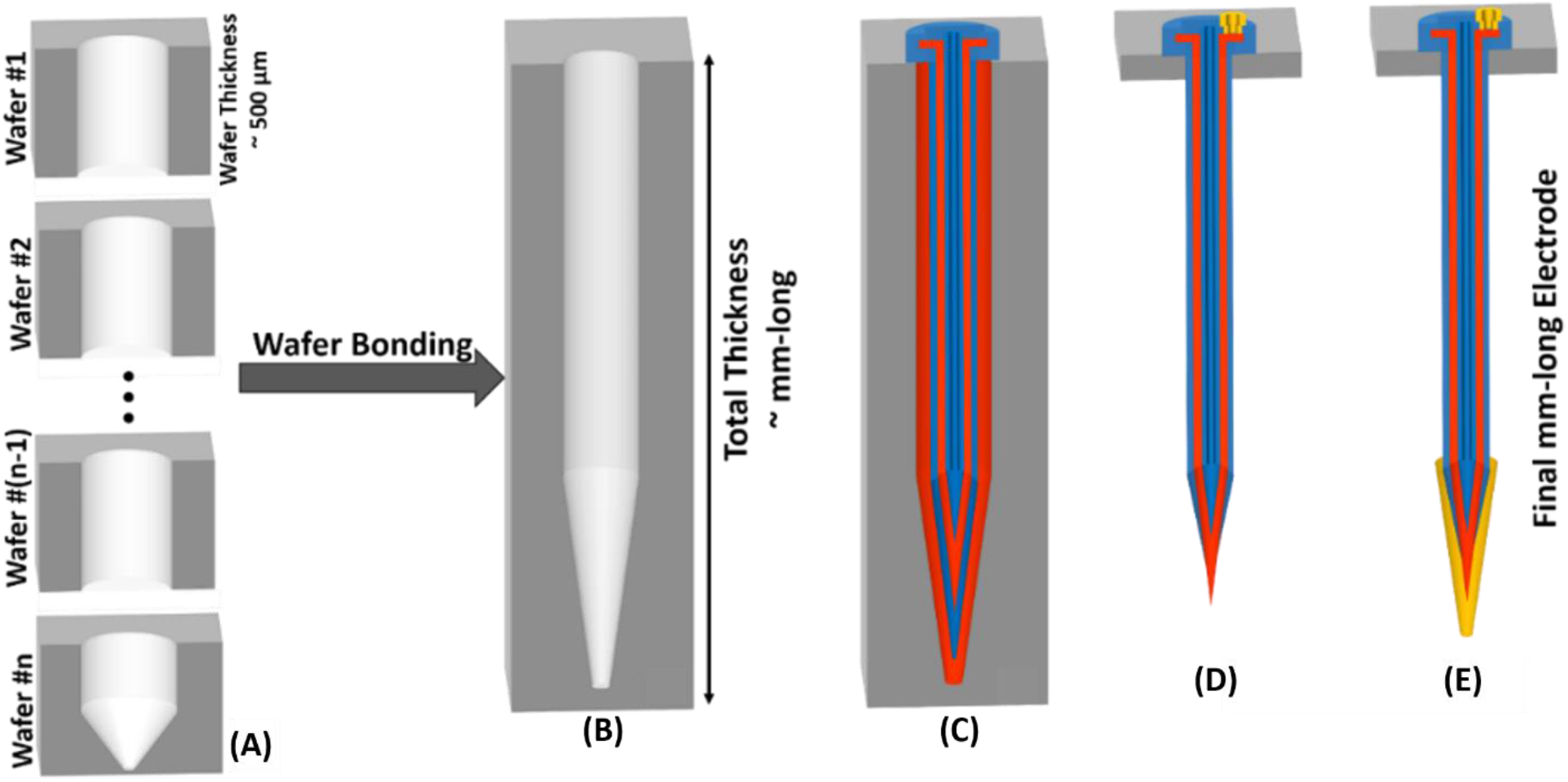
Fabrication of millimeter-long needles: (A) Process starts with preparing multiple separate wafers containing through-wafer etched holes using ultra-deep high-aspect-ratio ramped recipe and one wafer etched using a custom DRIE recipe to utilize the DRIE lag effect to form the tip shape and sharpness. (B) Next, all the wafers are bonded to form a mm-deep hole, (C-E) then the holes will be refilled with the same fabrication process described in Fig. 3.1 to obtain mm-long electrode arrays.

Millimeter-long probe arrays fabricated using the bonding-refilling technique described above are shown in Fig. 6. In this process, four 500μm-thick silicon wafers are bonded using fusion bonding to obtain ~1.7 mm deep holes. The bonded stack of wafers is then processed using the fabrication technique described in Fig. 3. 1.2mm long probe arrays with various number of shanks, pitch and size are realized by etching the top three silicon wafers away and leaving the bottom wafer as the substrate. For longer needles, more wafers can be bonded. It is worth mentioning that the 10×10 array shown in Fig. 6(A) provides the same number of electrodes as Utah array on a footprint 16 times smaller than Utah array.

**Figure 6.**
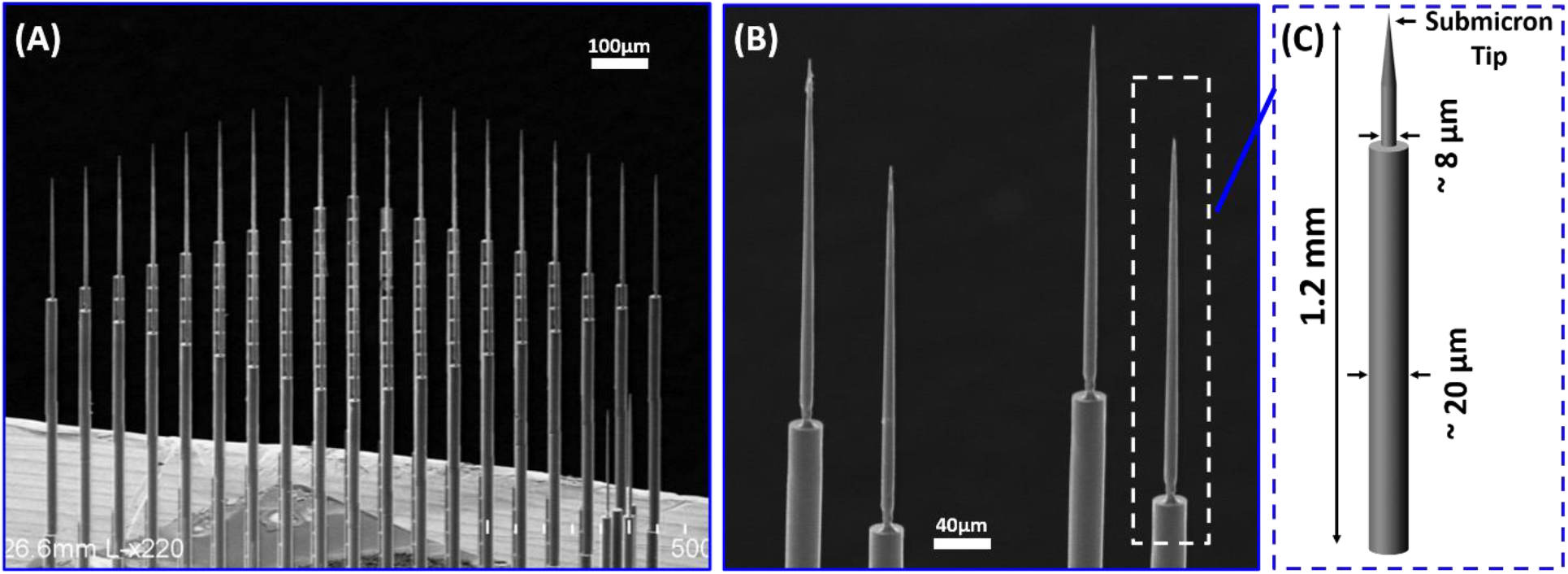
SEM images show the fabricated mm-long electrodes using the process described in Figure 5: (A) An array of 10×10 electrodes with 1.2 mm long shanks and a shank-to-shank pitch of 100μm are realized by bonding four 500μm-thick silicon wafers and etching away the top three wafers to keep the last wafer as a backbone. The 10×10 electrode array footprint is ~ 1mm×1mm which is 16 times smaller than the Utah electrode array (B) Close-up view of the conical-shaped upper section of electrodes (C) Electrode shanks are ~20 μm thick at the cylindrical base, ~8 μm thick at the tip conical base and submicron size at the tip.

### Overview of the Innovative Features

The broad goal of this research has been to develop a new class of extremely dense, high-count, and versatile electrode arrays for high-bandwidth high-resolution interfacing with irregular-shaped neural structures.

From a technology standpoint, the SEA technology design, fabrication, and structure are all novel and significantly different from current technologies. As explained in the previous sections, many kinds of multi-channel electrodes are available today, utilizing different fabrication technologies. One can categorize these fabrication technologies into two broad classes. One utilizes assembly and manipulation of individual electrodes (such as carbon fibers or microwire bundles), and the other relies on planar microfabrication technologies, such as those used for semiconductor electronics fabrication. Our approach allows us to combine the best features of both of these approaches. We can produce a very dense and high-count array of fine and slender insulated silicon needles with variable length and width, a tapered cross-sectional shape along their length, and sharp tips.

Fig. 7 illustrates the innovative features of the micro-electrodes that can be fabricated using the proposed technology. These features include:

1. Individual needles can have different lengths, and still reside side by side, and can be distributed in any arbitrary pattern. Needles as tall as several millimeters can be fabricated as needed in many applications.
2. Needles can have variable pitch/density, with needles capable of being spaced as close as 10μm.
3. Needles can have different shank diameters, which helps in engineering the stiffness of each shank. Shanks as small as a few microns in diameter can be fabricated. Obviously, there is a tradeoff between stiffness and flexibility, which can be tailored for different applications by the user.
4. Several needles with extremely small diameters can be fabricated in even closer proximity than typical needles and form a cluster, like a tetrode, to monitor neural activity in depth within the same column of tissue.

**Figure 7.**
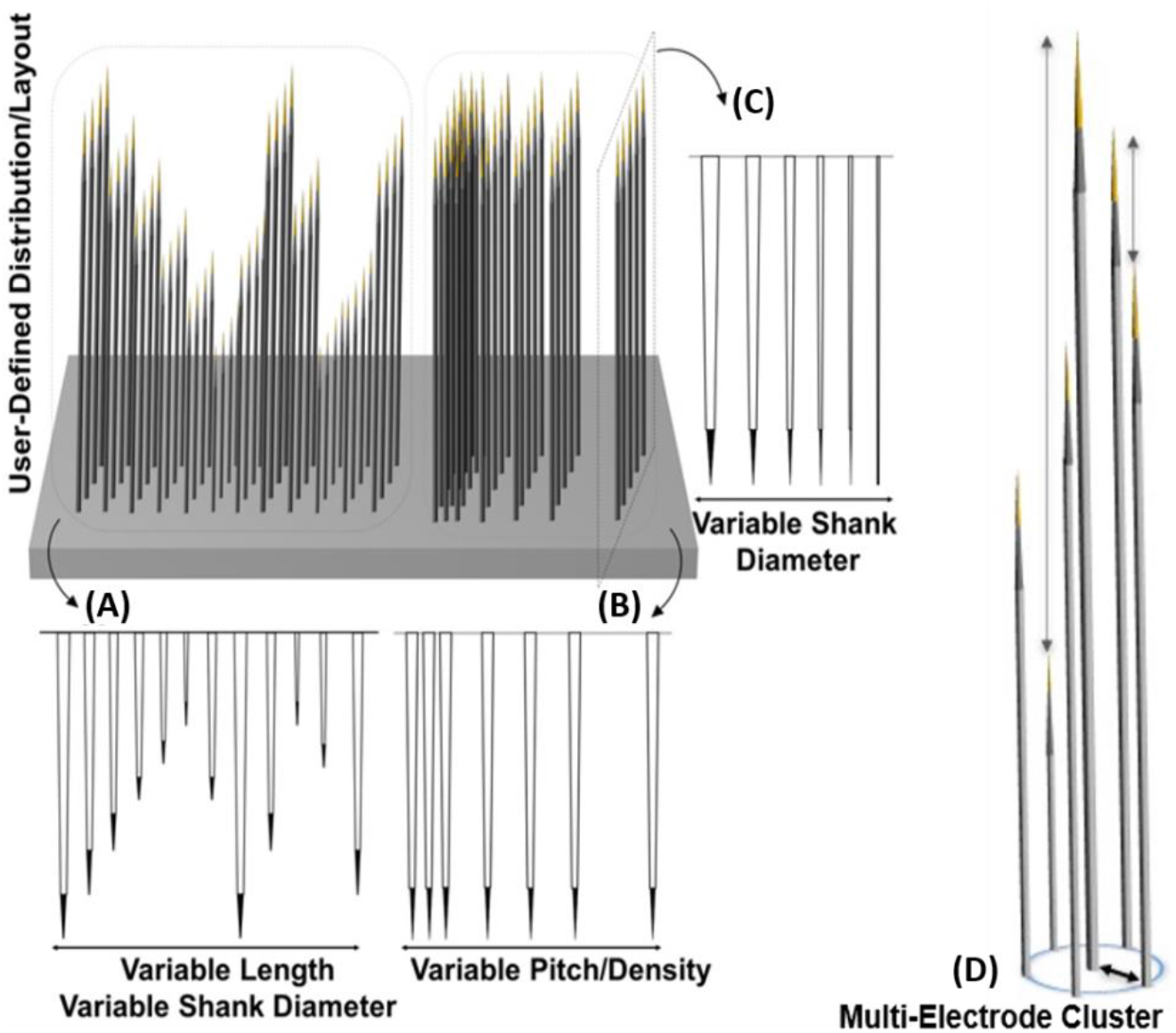
Innovative features of the proposed micro-needles: (A) Individual needles can have different lengths, and still reside side by side, can be distributed in any arbitrary pattern. Needles can be as tall as several millimeters. (B) Needles can have variable pitch/density, capable of being spaced as close as 10 μm. (C) Needles can have different shank diameters, to allow design of the stiffness of each shank. Shanks with diameters as small as a few microns can be fabricated. (D) A cluster of closely spaced needles with different heights will allow depth recording.

In the following sections, we will discuss the details of the unique features of the developed SEA technology.

### Probe Shank Conical Shape and Tip Sharpness

The conical shape of the shanks and tips shown above is naturally obtained through a special feature of the DRIE etch step called DRIE lag effect. DRIE etch rate is inversely proportional to the aspect ratio of the hole/trench, in other words, holes/trenches with higher ratio of depth to width are etched slower [29–35]. The variable etch rate at different depths will result in the conical shape of the shank, producing a sharp tip which improves probe insertion into the tissue. Fig. 8(A-C) shows three sets of electrode arrays fabricated with different starting hole sizes (15, 25, and 35 μm) resulting in shanks with different sizes and conical shape. Fig. 8(D-E) shows SEM images of electrodes with various tips size and shape which is controlled by changing the opening hole diameter. As shown in Fig. 8, holes with smaller diameters result in tips with smaller size and smaller opening angle and therefore increased sharpness.

**Figure 8.**
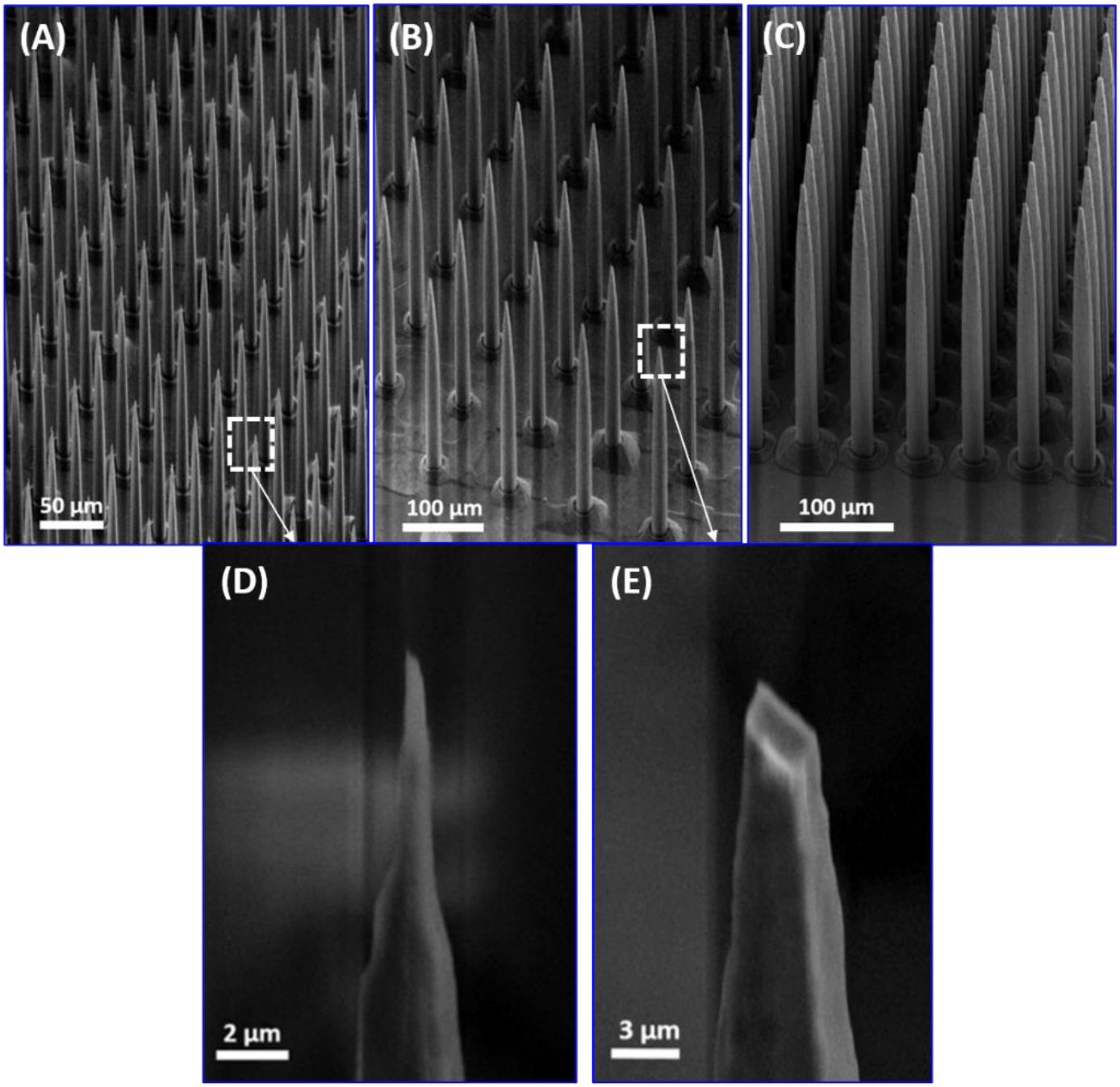
Fabricated electrode arrays using starting holes with different diameters: (A) ~10 μm (B) ~20 μm (C) ~30 μm thick probe shanks resulted from 15 μm, 25 μm and 35 μm hole diameter, respectively. SEM images of the submicron size electrode tips: (D) 10 μm thick, (E) 20 μm thick electrode shank, sharp tips and small probe shank size reduces the tissue damage during the implantation and post-implantation, improves the chronic stability and increases the array density.

### Varying Electrode Length, Size, Pitch and Distribution

There are several aspects of this fabrication technology that are critical for successful interfacing with neurons across multiple spatial latitudinal and longitudinal planes with high spatial resolution. First, the length of side-by-side electrodes can be varied using two methods:

1. DRIE lag approach: By changing the hole diameters in the layout, due to the DRIE aspect-ratio dependent etching (ARDE) and DRIE lag effect in holes with various depth, and electrodes with various lengths as shown in Fig. 9 can be produced.
2. Wafer bonding approach: By engineering the mask layouts used for patterning the holes in each wafer, the length of side-by-side electrodes within a die can be controlled.

**Figure 9.**
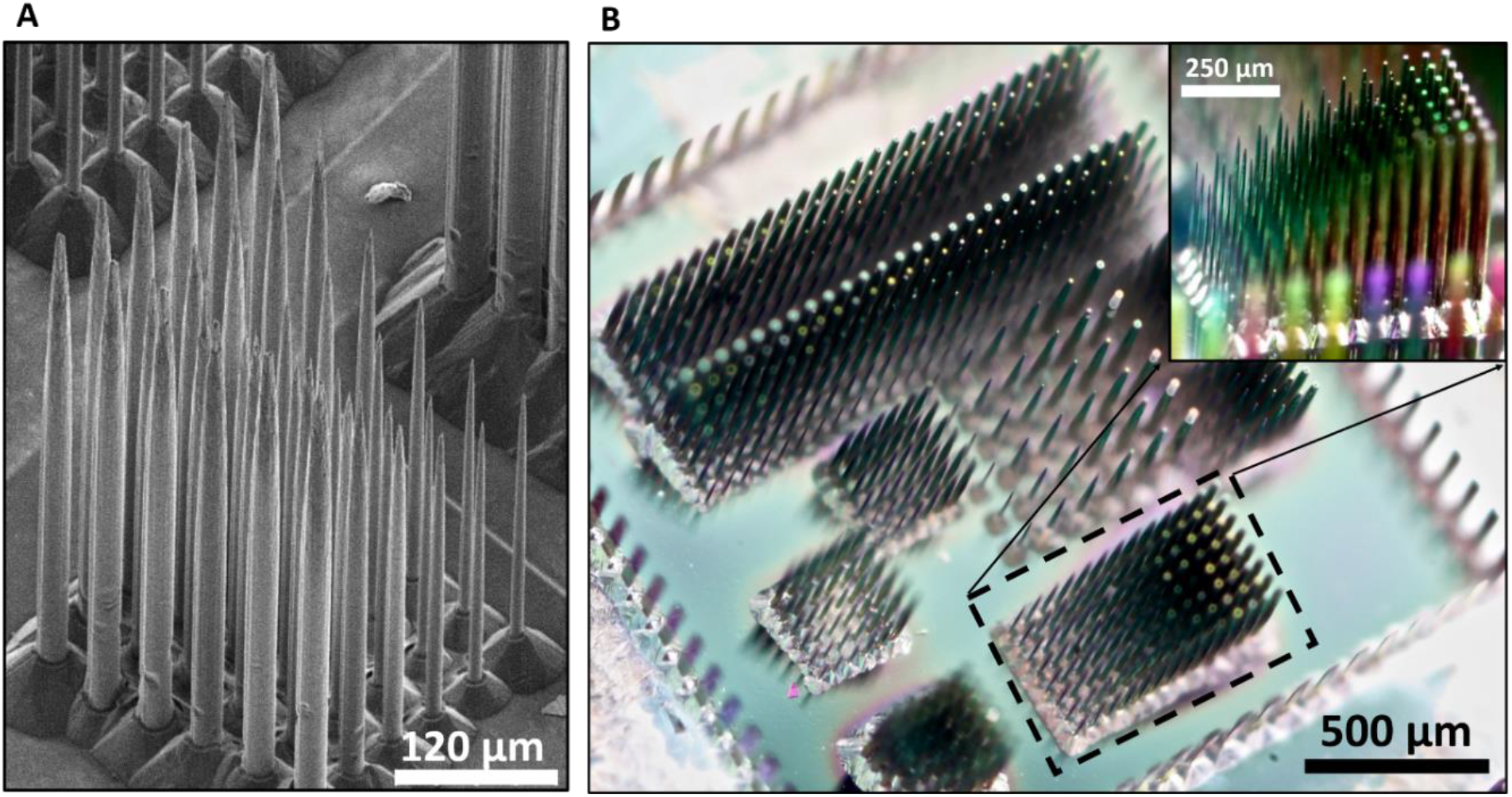
Arrays with various electrode length, thickness, pitch and distribution is realized using the refilling technology. These features enable recording and stimulation of larger number of neurons in the implanted tissue volume using minimum number of electrodes resulting in less tissue damage and improved chronic stability. (A) SEM image of fabricated side-by-side electrodes with varying length and diameter. (B) Optical image of an array with an arbitrary distribution of electrodes with various length, shank diameter and pitch.

In the first technique (DRIE lag approach), reducing the holes opening size results in reduced transport of the DRIE process etchant agents and ion bombardment at the bottom of the features as the aspect-ratio increases throughout the deep etching process. This will cause reduced etch rate, tapering and convergence of the holes sidewalls, and eventually termination of the etch process. We have utilized this self-terminating process to obtain side-by-side electrodes with different heights (lengths) using holes with various opening diameters in a single process as shown in Figure 5.4. Using this technique, the maximum variation of length is limited to 500 μm for reasonable hole diameters (<35 μm). To extend the range of electrodes length, another technique based on wafer bonding is developed as described in the following.

In the second technique, electrodes length is dependent on the hole layout design in each substrate of the bonded stack. In other words, larger holes (20-35 μm) are replaced by small holes (<10 μm) to form a conical tapered hole instead of thru-wafer holes with straight sidewalls. Depending on the substrates thickness and the relative position of hole sizes reduction along the bonded substrates stack, the final depth of the holes and therefore final electrode shank length is determined.

The electrode pitch can be locally modified within the array, in other words the density of recording sites can be customized based on the application by changing the layout. Studies have shown that the electrode size plays an important role in determining tissue damage and chronic stability of the implanted array [36]. Any arbitrary distribution of electrodes with various length, shank diameter and pitch across the array is obtained. Optical and SEM images show the fabricated array with all the features mentioned above in Fig. 9 Employing the mentioned features (varying side-by-side electrode length, pitch, and distribution) will enable 3D access of neurons with extremely high spatial resolution. In particular the varying length feature is critical in implantation into the brain convoluted surface and simultaneous recording/stimulation of neurons at various depth of tissue.

### SEA Characterization: Impedance Spectroscopy, Mechanical Robustness and Acute *in vivo* Results

Electrochemical Impedance Spectroscopy (EIS) is used to characterize the recording sites electrical properties. Fig. 10(A) shows the setup used to measure a 3×3 SEA recording sites impedance. Fig. 10(B) shows the impedance magnitude and phase values for one of the recording sites at frequencies between 10 Hz and 50 kHz. Fig. 10(C) shows the impedance magnitude and phase values for all 9 recording sites measured at 1 kHz.

**Figure 10.**
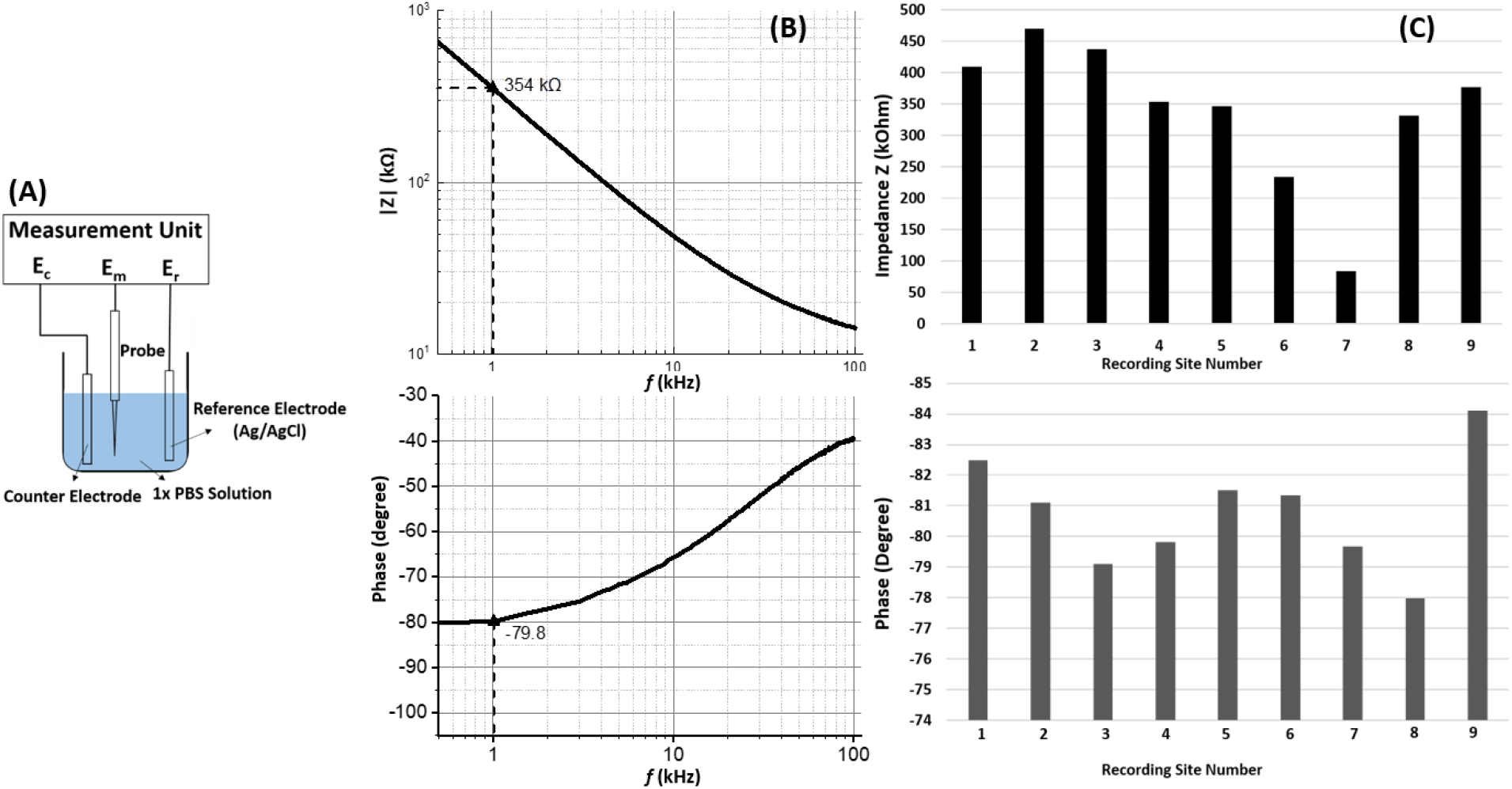
Electrode recording site impedance spectroscopy setup and measurement results. (A) EIS setup consists of a counter electrode (stainless steel), reference electrode (Ag/AgCl), 1x PBS solution, a measurement unit and the electrode. (B) Impedance measurement results (amplitude and phase) of one of the recording sites which is measured to be 410 kΩ at 1 kHz. (C) EIS measurement results (impedance magnitude and phase) for 9 electrodes recording sites measured at 1 kHz.

### Mechanical Properties of the Electrodes

Mechanical robustness issue and in particular buckling and breakage are the main failure modes for penetrating neural probes during insertion and post-implantation. We have studied the effects of electrode geometry on tissue insertion using the linear buckling analysis in COMSOL Multiphysics. Fig. 11 shows the 3D and 2D images of the simulated electrode geometry and materials in these studies which are chosen to represent the fabricated electrodes described For the buckling load analysis, a fixed-hinged boundary condition is chosen, meaning that the shank base is fixed and the tip has a hinged boundary condition in the shank axial direction with no lateral displacement as illustrated in Fig. 12. Table 1 shows the material properties used in these simulations.

**Figure 11.**
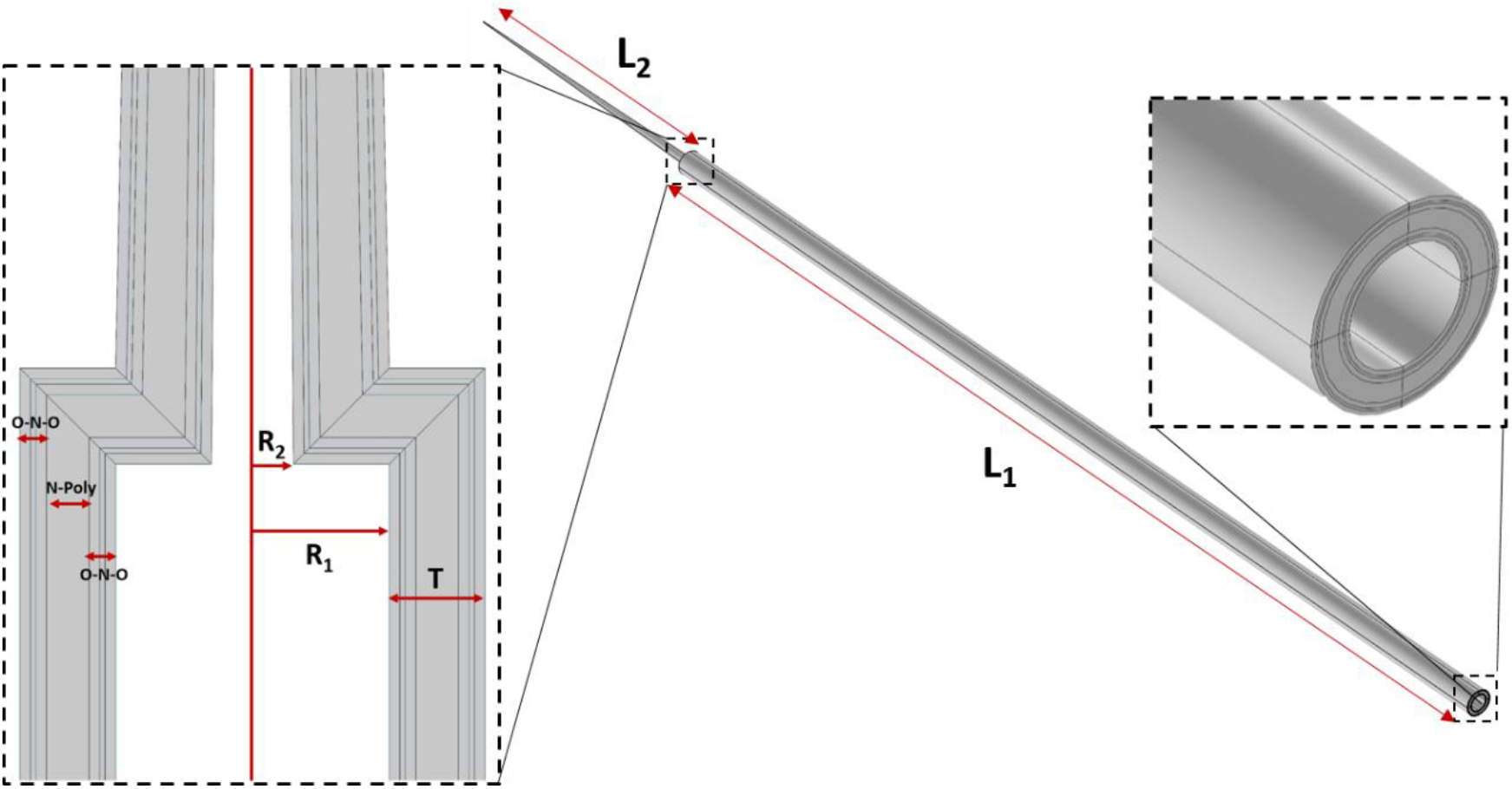
2D and 3D structure of modeled electrode geometry and materials in COMSOL Multiphysics.

**Figure 12.**
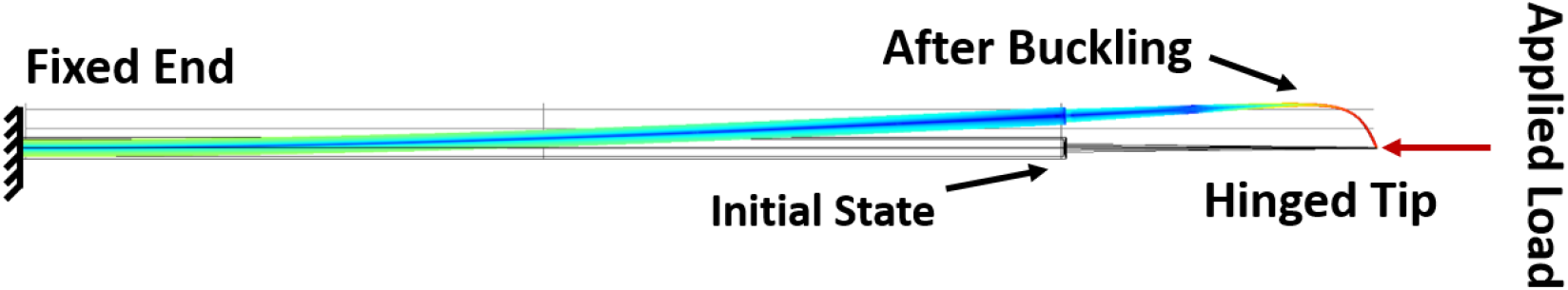
Boundary conditions of linear buckling load analysis: Electrode shank base is fixed while the electrode tip has a hinged boundary condition.

**Table 1.**
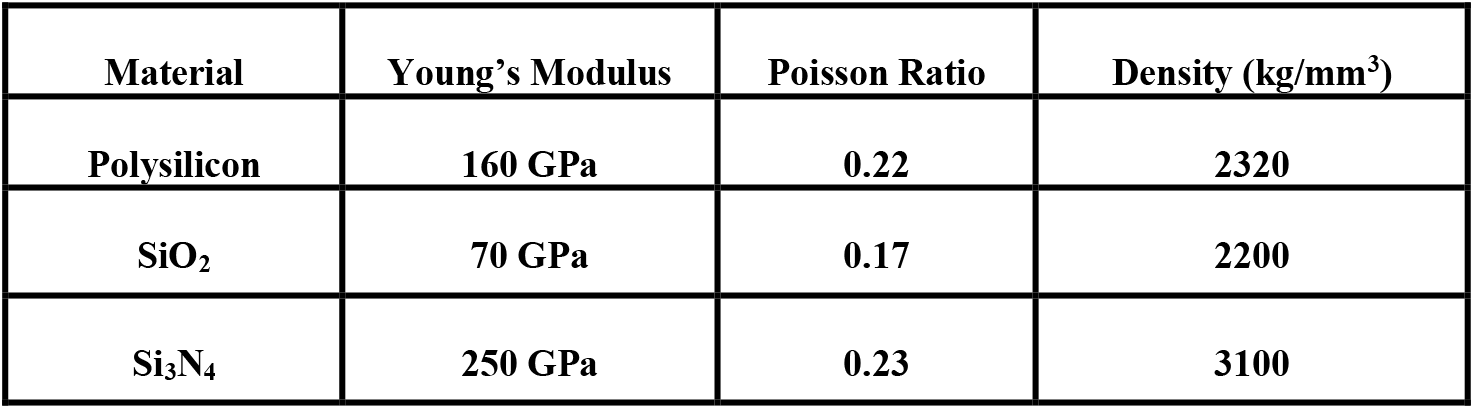
Material properties used in the buckling load analysis simulations.

The electrode length is one of the key parameters that determines the shank critical buckling force, meaning that longer electrodes exhibit smaller critical buckling force. Critical buckling load modeling and simulations is used for the dimensions shown in Table 2 to obtain a safe range of electrode length that ensure successful rat brain pial penetration without buckling and breakage.

**Table 2.** The electrode geometry parameters used in the buckling load simulations.

|  |  |
| --- | --- |
| Cylindrical Base Length ($L_1$) | Swept [1-12 mm] |
| Conical Tip Length ($L_2$) | Swept [0.15-0.25 mm] |
| Shank Thickness (T) | 4 $\mu\text{m}$ |
| Tip Radius ($R_3$) | 0.25 $\mu\text{m}$ |
| Base Inner Diameter ($R_1$) | 7 $\mu\text{m}$ |
| Tip Base Diameter ($R_2$) | 5 $\mu\text{m}$ |

The closest probe used in this work to our structure was a tungsten electrode that was manually electro-sharpened with an opening angle of 4° and shank diameter of 50 μm. This probe exhibited a penetration force of ~0.62 mN [38]. Therefore, we assume the required pia penetration forces to be 0.62 mN, although we expect it to be smaller for our probe as the shank diameter is smaller. This is a very conservative but safe assumption for determining the penetration forces and critical buckling load values. Fig. 13 shows simulations results of critical buckling load of electrodes with R_1_ = 7 μm, R_2_ = 5 μm and various L_1_ and L_2_ values. As shown, the maximum length to obtain critical buckling load larger than 0.62 mN is limited to 4mm. Fig. 14 visually demonstrates the effect of L_1_ on the electrode buckling shape and von Mises stress distribution along the shank. As shown, in electrodes with smaller L_1_, buckling mainly occurs at the conical part since this part has a smaller diameter, while for longer electrodes, buckling occurs along the longer cylindrical part of the shank.

**Figure 13.**
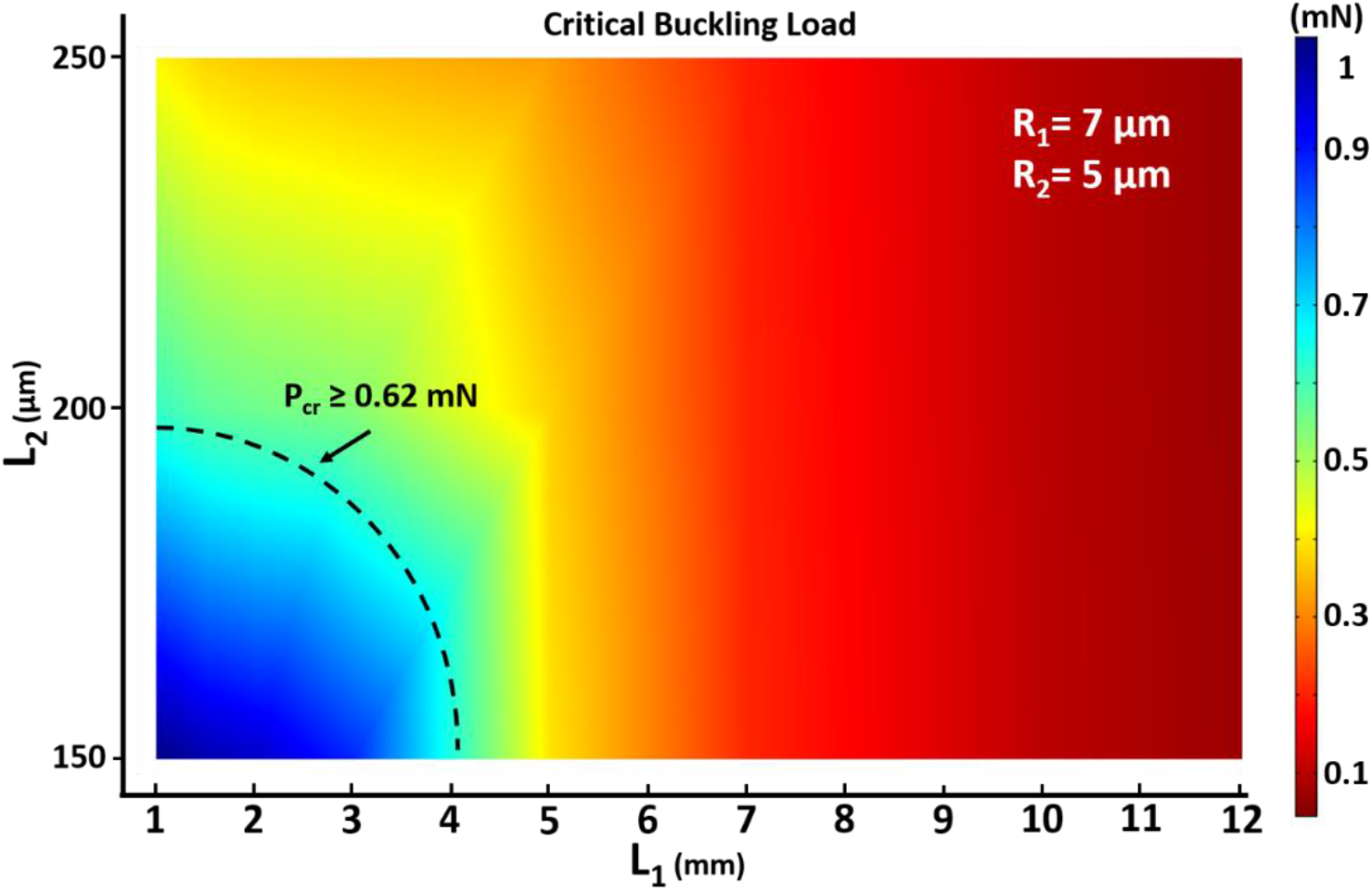
Simulation results of critical buckling load of electrodes with R_1_ = 7 μm, R_2_ = 5 μm and various L_1_ and L_2_ values. As shown, the maximum length to obtain critical buckling load larger than 0.62 mN is limited to 4mm.

**Figure 14.**
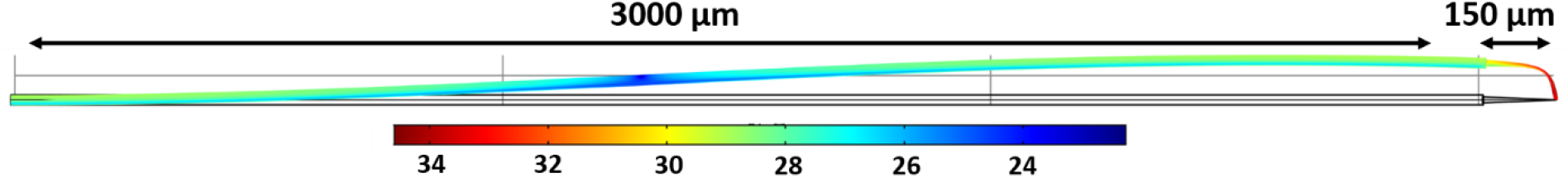
Effect of electrode length on the buckling shape and von Mises stress distribution along the shank. In shorter electrodes buckling mainly occurs along the conical part of the tip while in longer electrodes, buckling is more evident along the cylindrical part of the shank.

Mechanical robustness and flexibility of the electrodes is tested by applying a bending force at the tip of the electrodes using a micromanipulator probe tip under the probe station microscope. Fig. 15(A-B) shows the electrode before and during application of the bending force at the electrode tips. As shown, the electrode can bend for ~28° with ~115 μm lateral displacement at the tip before breakage. Fig. 15(C) shows that breakage ultimately occurs at the junction of the conical tip and cylindrical base as expected.

**Figure 15.**
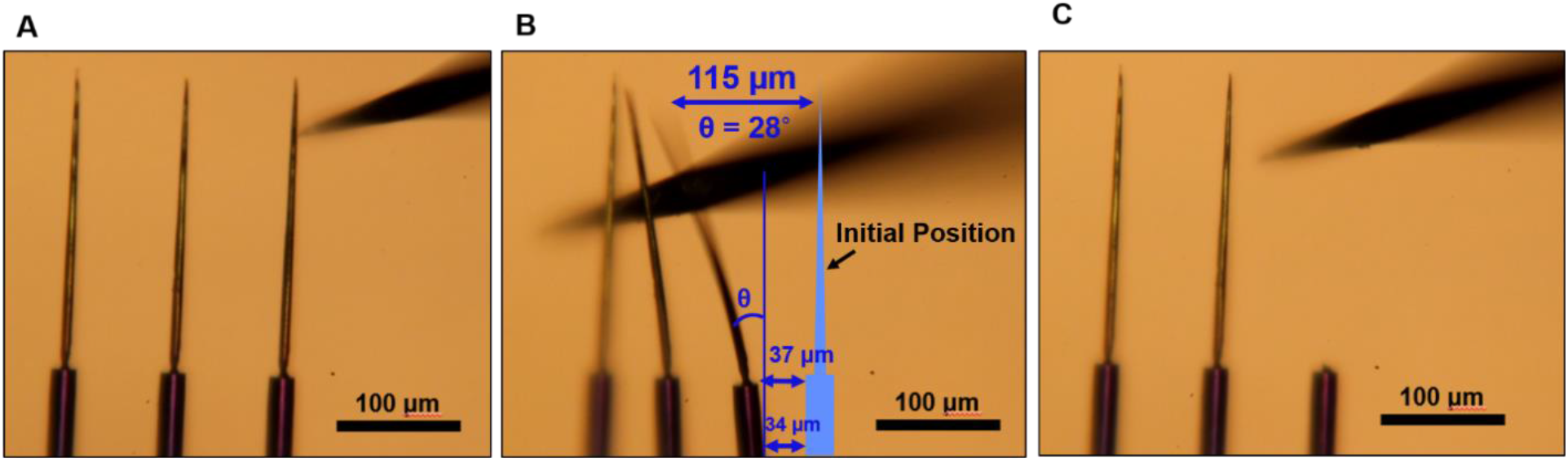
Mechanical characterization of electrodes using a bending test: Robustness and flexibility of electrodes is tested by applying a bending force at the electrode tips using probe station tips. (A) Electrodes before applying the force, (B) Electrodes maximum bending and lateral displacement right before breakage, the electrode tip is bent ~ 28° and displaced ~ 115 μm laterally before breakage occurs. (C) The breakage occurs at the junction of the tip conical part and cylindrical base (the most right electrode).

Penetration of an electrode through the rat brain pia mater is studied using finite-element analysis. The model consists of three components, an electrode, a thin elastic layer representing the pia mater membrane and an elastic bulk material representing the brain tissue beneath the pia. The electrode geometry parameters obtained in previous sections are used in this simulation as shown in Table 3.

**Table 3.** Electrode geometry parameters used in the pia mater penetration simulations.

|  |  |
| --- | --- |
| **Cylindrical Base Length ($L_1$)** | **1 mm** |
| **Conical Tip Length ($L_2$)** | **0.15 mm** |
| **Shank Thickness ($T$)** | **$4 \mu\text{m}$** |
| **Tip Radius ($R_3$)** | **$0.3 \mu\text{m}$** |
| **Base Inner Diameter ($R_1$)** | **$9 \mu\text{m}$** |
| **Tip Base Diameter ($R_2$)** | **$5 \mu\text{m}$** |

The pia layer 3D model is a cylinder with a thickness of 10 μm and diameter of 1 cm. Similarly the brain tissue has a cylindrical structure with height and diameter of 5 mm and 1 cm respectively. Table 4 presents the material properties used in the brain model to accurately match the rat brain structure [39, 41–44]. To examine whether the electrode can successfully penetrate through the pia layer without buckling, we applied a force at the other end of the electrode and simulated the von Mises stress distribution through the pia layer volume. The applied force is chosen to be below the electrode critical buckling force. Based on the linear buckling simulation results obtained in previous sections, the critical buckling load of the electrode used in this simulation is around 665 μN. Therefore, we applied a load much smaller than the critical buckling load (100 μN) to the electrode to prevent buckling. Fig. 16 shows the von Mises stress distribution in the 3D model of the electrode-pia-tissue structure. These simulations show that the pia layer stress reaches as high as 120 MPa which is much larger than the required stress needed for penetration through pia (40 MPa) as reported in [38, 40, 45]. Therefore, this model predicts the designed electrode is capable of penetrating the rat pial membrane without buckling. Also, the small volumetric stress concentration is a sign of minimal dimpling and tissue stress which demonstrate the importance of tip size and sharpness.

**Table 4.** Material properties used in the brain model to accurately match the rat brain structure [39, 41–44].

| Material | Young's Modulus | Poisson Ratio | Density (kg/mm<sup>3</sup>) |
| --- | --- | --- | --- |
| Pia Mater | 15.8 MPa | 0.45 | 1130 |
| Brain Tissue | 30 kPa | 0.45 | 1050 |

**Figure 16.**
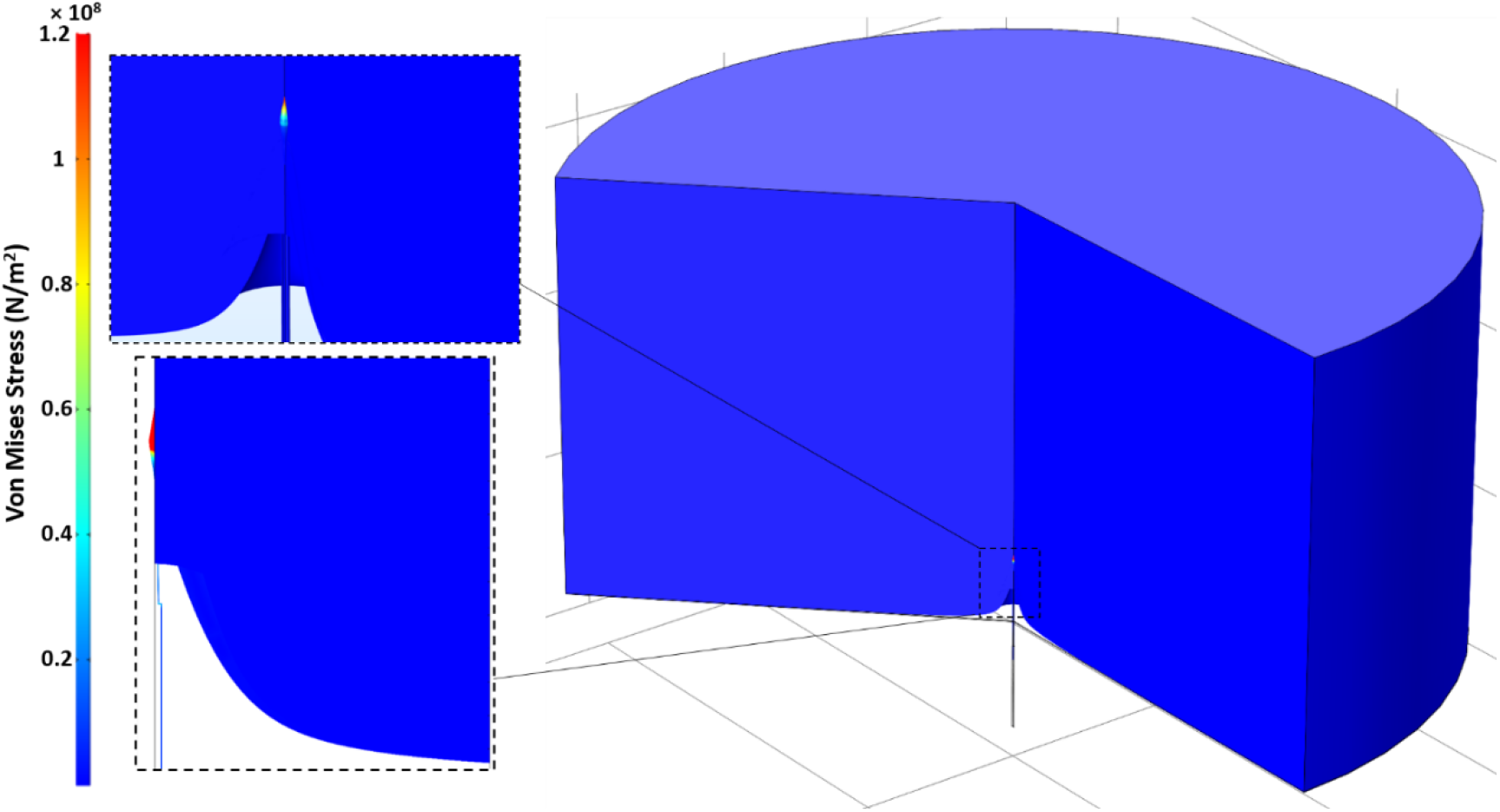
Von Mises stress distribution in the 3D model of the electrode-pia-tissue structure. By applying a 0.1 mN load the pia layer stress reaches as high as 120 MPa which is much larger than the required stress needed for penetration through pia (40 MPa). Therefore, this model predicts the designed electrode is capable of penetrating the rat pia membrane without buckling, also, the small volumetric stress concentration is a sign of minimal dimpling and tissue stress which demonstrated the importance of tip size and sharpness.

### Acute *in vivo* Studies

The first step to perform acute studies is to make electrical interconnections between the electrodes and the external recording equipment. We used arrays with low electrode-count (<100 electrodes) and large pitch size (100μm-500μm) and millimeter-long shanks (1mm-1.5mm) for acute studies. Upon incorporating the interconnections, the device is ready for acute *in vivo* studies as shown in Fig. 17(A). We have used the electrodes with lower site impedances for *in vivo* recording. Acute studies are performed on mice/rats under anesthesia. The SEA arrays manufactured using the microfabrication process described in Fig. 5. 2×2 and 3×3 arrays of millimeter-long electrodes were implanted in a rat brain under anesthesia. The site impedance of the probes (the sites were located just below the tip), was in the range of a few hundred kΩ at 1 kHz as shown in Fig. 10. The sharpness of the probes ensured an extremely straightforward insertion as shown in Fig. 17(B). After recordings, the probes consistently remained fully intact as shown in Fig. 17(C), indicating their stiffness despite their narrow, small footprint. Recordings under isoflurane anesthesia showed characteristic slow field potentials, along with occasional epochs of 11 Hz oscillations shown in Fig. 17(C) and Fig. 17(D) that have been recently reported in recordings under isoflurane [37]. Thus, we have already performed the first series of implants to establish that the probes are viable, easy to insert, resistant to breakage, and able to record electrical signals.

**Figure 17.**
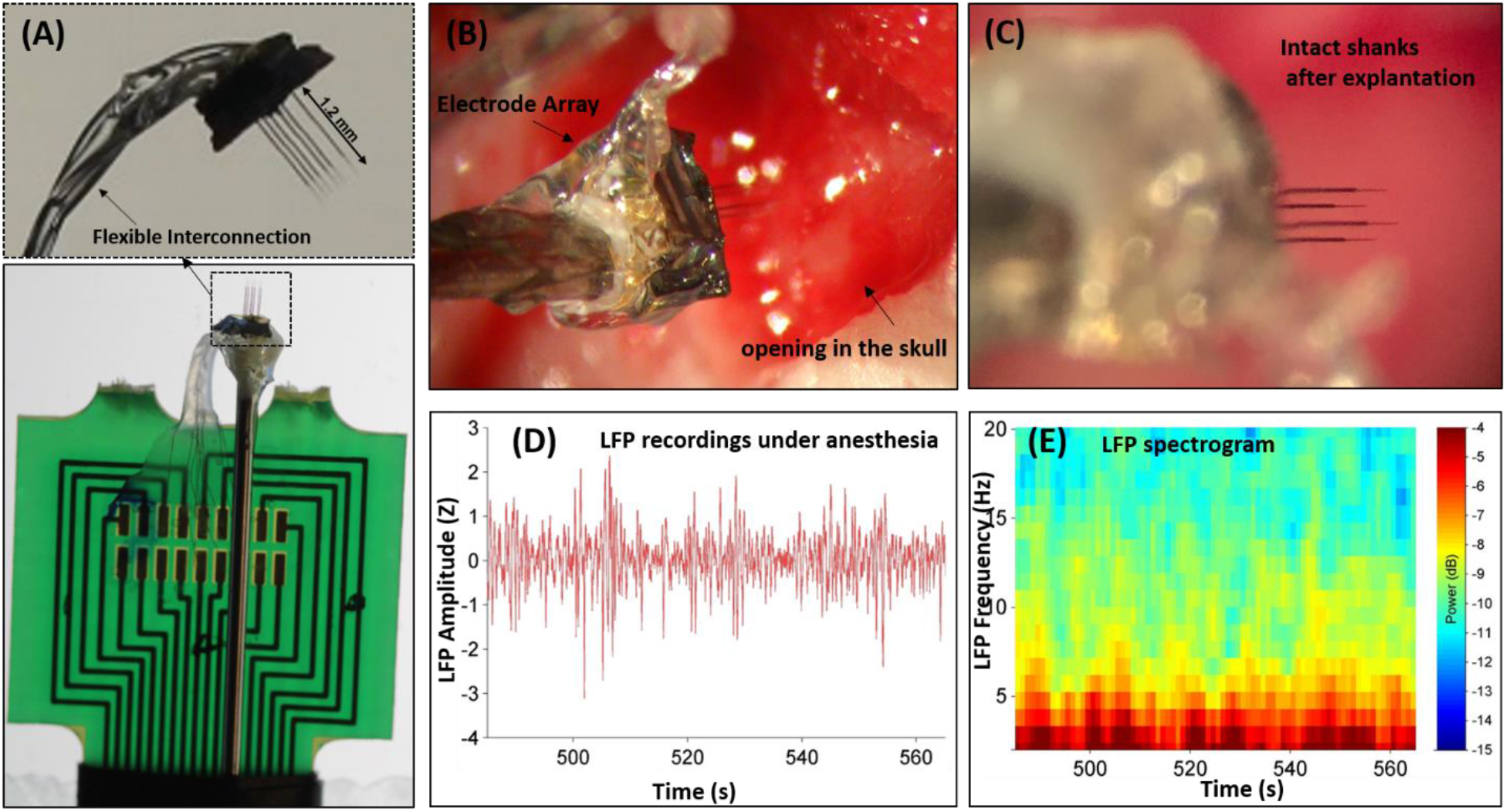
Acute recordings using the SEA array *in vivo*: (A) Electrodes pads are connected to a PC board using wire bonding. To insulate the wires and enhance the flexibility and robustness, an epoxy will be applied to the wires to form a highly flexible insulated wire-bundle. (B) An image of a prototype SEA array during implantation into rodent brain. The array was able to seamlessly penetrate brain tissue thanks to the sharp needle design of the probes. (C) Image of SEA array after removal from the brain showing fully intact probes. (D) LFP recordings in a rat barrel field cortex under isoflurane anesthesia recorded on one of the channels of the SEA arrays. (E) Spectrogram of the LFP recordings showing characteristic slow oscillations and ~ 11 Hz epochs that are both hallmarks of LFP under isoflurane anesthesia, demonstrating the ability of these probes to record high-quality neural signals.

## Conclusion

This paper presented design and development of a new silicon-based fabrication technology for producing high-density, large-count, 3D arrays of extremely fine electrodes with user-defined length, width, shape and tip profile. Simultaneous recordings from, and stimulation of, a large number of neurons with high spatiotemporal resolution across multiple spatial planes is crucial to decipher complex neural circuits with causal single cell precision. Near-ideal neural probe features are described as a guideline to design and develop a new electrode array technology. To overcome the shortcomings and issues of previously reported penetrating neural probe technologies, a fabrication process based on refilling deep ultra-high aspect-ratio holes in a silicon substrate with deposited layers followed by etching away the support substrate to leave thousands and eventually millions of needles, is developed. Electrode arrays with various design parameters are fabricated to demonstrate the capabilities of the proposed technology. This includes arrays with different electrode count (4 - 5184), pitch (50μm-500μm), electrode length (200μm-1.2mm), diameter (10μm-30μm) and arbitrary distribution of electrodes with varying length, pitch, diameter across the array. These results demonstrate our technology capabilities to make high-density large-scale arrays with true 3D spatial coverage which are characteristics of the near-ideal probe. Functionalities of the developed SEA array is tested by acute *in vivo* recordings in a rat barrel field cortex using 2×2 and 3×3 arrays. The array was able to seamlessly penetrate the pia mater and tissue with no tissue dimpling or electrode buckling thanks to the sharp needle design of the probes. Local field potentials (LFP) recordings under isoflurane anesthesia showed characteristic slow field potentials, along with occasional epochs of 11 Hz oscillations.

We believe the work presented in this paper has provided solutions to some of the many challenges that prohibit realization of neural interfaces for therapeutics and brain-machine interface applications, however, further research is needed to address the unresolved problems. These include: Fabrication of electrodes with smaller cross-sectional size to alleviate the tissue inflammatory response. Modification of the recording site material to improve the recording site impedance and reliability of the sites to obtain chronically stable recordings of single units. Alternative dielectric and conductive materials for improved impedance and stability of electrodes *in vivo.* Examples of alternative dielectrics include Al2O3 or SiC deposited using Atomic Layer Deposition (ALD) or Chemical Vapor Deposition (CVD) processes, respectively. Pt, Ir and/or electrodeposited PEDOT is suggested as the recoding site material to improve the impedance and recording signal to noise ratio. Modification of fabrication technology to realize other recording/stimulation modalities such as optical and chemical. Integration of all these modalities (electrophysiological, optical and chemical) in a single probe can provide neuroscientists a great tool to advance our understanding of the brain.

